# BioCompute Objects to communicate a viral detection pipeline with potential for use in a regulatory environment

**DOI:** 10.1101/2021.10.19.465010

**Authors:** Naila Gulzar, Jonathon Keeney, Jack B. Baker, Ondrej Klempir, Geoffrey Hannigan, Danny A. Bitton, Julia M Maritz, Charles Hadley S. King, Janisha A. Patel, Paul Duncan, Raja Mazumder

## Abstract

The volume of nucleic acid sequence data has exploded in recent years, and with it, the challenge of finding and transforming relevant data into meaningful information. Processing the abundance of data can require a dynamic ecosystem of customized tools. As analysis pipelines become more complex, there is an increased difficulty in communicating analysis details in a way that is understandable yet of sufficient detail to make informed decisions about results or repeat the analysis. This may be of particular interest to institutions and private companies that need to communicate complex computations in a regulatory environment. To meet this need for standard reporting, the open source BioCompute framework was developed as a standardized mechanism for communicating the details of an analysis in a concise and organized way, and other tools and interfaces were subsequently developed according to the standard. The goal of BioCompute is to streamline the process of communicating computational analyses. Reports that conform to the BioCompute standard are called BioCompute Objects (BCOs). Here, a comprehensive suite of BCOs is presented, representing interconnected elements of a computation that is modeled after those that might be found in a regulatory submission, but which can be shared publicly. Because BCOs are human and machine readable, they can be displayed in customized ways to further improve their utility, and an example of a collapsible format is shown. The work presented here serves as a real world implementation that imitates actual submissions, providing concrete examples. As an example, a pipeline designed to identify viral contaminants in biological manufacturing, such as for vaccines, is developed and rigorously tested to establish a rate of false positive detection, and is described in a BCO report. That pipeline relies on a specially curated database for alignment, and a set of synthetic reads for testing, both of which are also descriptively packaged in their own BCOs. All of the sufficiently complex processes associated with this analysis are therefore represented as BCOs that can be cross-referenced, demonstrating the modularity of BCOs, their ability to organize tremendous complexity, and their use in a lifelike regulatory environment.

## Introduction

### Virus Detection using HTS and challenges

High-throughput sequencing (HTS) has become an indispensable approach for the identification and characterization of pathogens. The application of HTS is remarkable in the field of detection of known and unknown viral nucleotide sequences that may be present at low amounts in complex biological samples [1] [2]. The potential of HTS to produce millions of short nucleotide sequences from a sample, without prior knowledge about the viral genomes, in a time-efficient and cost-effective manner, has led to the discoveries of novel viruses [3] [4] [5].

HTS is a powerful technology for the screening of vaccines, particularly the live attenuated vaccines, to maintain safety standards [6] [7], and for investigation of the outbreak of novel viruses [8] [9] [10]. However, HTS poses many challenges due to the presence of contaminations incorporated at the sample preparation step of HTS, as well as scope and curation of the reference database that is queried to map the HTS reads [11] [12]. Previous studies have shown contaminations in the form of laboratory components containing viral nucleotide sequences [1] [13] [14]. In addition, simple environmental contamination with human and various animal sequences in reagents and water may confound routine analyses against reference databases populated with normal endogenous retro element sequences or with references possibly containing viral analogs of normal cellular genes (such as oncogenes, ubiquitin, widely used vector components, and others). Therefore, rigorous approaches are required to establish the association of the novel viral genomes with the diseases. Great care must be taken in the development of an analysis pipeline, and caution in interpretation of the results, which is only possible with a very clear communication of the pipeline and associated tools. Given the complexity of the samples, the computations, the database, and the interpretations, an accurate and standardized means of communicating the computational parts of a study can facilitate understanding by scientific and regulatory stakeholders. The present study is therefore a very good example of a use case for regulatory submissions.

BioCompute is an open source project with roots in community discussions as far back as 2012. In 2014, the United States Food and Drug Administration (FDA) convened an internal meeting with its genome sequencing experts to discuss the growing problem of reviewer burden in evaluating submissions that contained a gene sequencing analysis, which were often delayed by months for lack of clear communication. Since then, several members of academia and the private sector have participated in shaping the project[15, 16], which became an approved IEEE standard in January of 2020[17], and which is now accepted by three Centers at the FDA for four drug applications that incorporate bioinformatics computations[18].

BioCompute is contrasted with workflow management languages like Common Workflow Language[19], Snakemake[20], Workflow Description Language[21], Nextflow[22], and Ruffus[23], in that these languages are specifically adapted to the computational management of a workflow, and are not well adapted to reporting in a regulatory environment. A BCO report does capture information about the execution environment (such as script drivers, specific package dependencies, and other system variables or prerequisites), but has a heavy emphasis on the metadata associated with a computational pipeline. A BCO is a human and machine readable report (written in JavaScript Object Notation) that is partitioned into conceptually related sections (e.g. the Parametric Domain for parameters, the IO Domain for input and output, etc.). A metadata record makes possible a variety of uses, such as a database of pipelines that is searchable by purpose, data type, or anything else, a manifest of incoming files, a peer review and certification of specific pipelines, or a quick verification test that allows a reviewer to determine if the results of the pipeline fall within its allowable error[24]. A BCO can also be paired with a workflow language to enable portability of execution, or with a Research Object[25] to package the full content of an experiment, and indeed active collaborations are currently underway to write documentation for joint implementation, as well as examples.

Because there is no other similar official method to communicate computational algorithms to the FDA, the alternative, and heretofore dominant way of communicating computational analyses to regulatory agencies has been ad hoc (personal communication). The growing importance of computational analyses, the nebulous character of the components of those analyses, and the rapidly changing nature of this relatively new field with new analyses regularly being developed mean increased organizational burden for many regulatory reviewers, and intensify the need for a streamlined communication vehicle. This is compounded by the rapid increase in regulatory filings that include complex computational analyses for which there may be comparatively few experts, making it more difficult to communicate novel strategies. While the standard is well documented and has a robust community and other efforts have focused on making the standard easier to work with[26, 27], the application of a standard may often present questions regarding implementation, and it is useful to have a real example of the way in which the standard is applied.

Here, an algorithm for adventitious virus detection in biological manufacturing processes using the HIVE platform, coupled with a curated reference viral database and synthetic reads for pipeline testing, is presented. Included is a comprehensive set of tools for such an analysis, including an instance of an open source, cloud-scalable platform for computations, a template pipeline for high sensitivity detection of viral reads in a variety of backgrounds, and a curated database of viral genomes. All of the elements are open source, and are provided as BioCompute Objects (BCOs) for clarity, which describes applicability and computational processes of the workflows, as well as the first published use of the user-described “Extension Domain,” used here in a way that is similar to the way in which supplementary information in publication might be used. In the case of the pipeline, a BCO is provided as a template (tBCO), using synthetic reads. By using this mock detection algorithm that is similar to a regulatory submission and its associated components, we demonstrate the utility of a BCO to coordinate several complex parts in an intuitive format that is easy to navigate and which reduces organizational burden on the receiver, in the hopes of improving efficiency of analysis communication.

### Algorithm for adventitious virus detection

The algorithm developed here uses the High-performance Integrated Virtual Environment (HIVE) bioinformatics platform[28] [29] for analysis of public datasets for detection of potential unexpected viruses and/or contaminants. HIVE is a cloud-based infrastructure with distributed storage and compute architecture, primarily meant for handling big data from HTS technologies. The initial codebase for HIVE was deployed in 2017 as an open source repository (https://github.com/FDA/fda-hive). The instance of HIVE used here is an adaptation of that project has been updated and refined, and can be accessed by the general public (see methods).

The volume of biologically meaningful data produced due to the analysis of extra-large datasets is projected to increase rapidly[30] [31] [32]. Therefore, methods and algorithms for deposition, storage and computations of these datasets need to be efficient, secure and have high levels of integrity. HIVE is designated as a FISMA-moderate equivalent (Federal Information Security Management Act) system [29] that provides secure access control and permissions for data sharing. The massively parallel distributed computing infrastructure of HIVE enables the distributed storage library and distributed computational powerhouse to connect seamlessly, creating a robust and flexible system by maintaining the storage and metadata databases on the same network [29]. The metadata related to extremely large files and the computations run in the HIVE system are stored in a metadata database and can be retrieved whenever needed. The resources available in HIVE include native algorithms, tools, and applications optimized for the HIVE architecture, as well as commonly used external tools adapted to operate within the HIVE infrastructure [28].

The viral detection pipeline was constructed using a specially curated version of the FDA’s Reference Viral Genomes Database[12] (RVDB), and tested through *in silico* read set generation, also carried out in HIVE. The pipeline uses HIVE tools such as the DNA *in silico* tool [28] for synthetic read generation, HIVE-Hexagon aligner[33], and the “alignment profiling tool,” HIVE-Heptagon[34]. These tools allow evaluation of the robustness of the detection settings and illustrate the interpretation of the final output for its ability to discriminate between potential non-viral versus unique and specific virus sequences.

As with many computational analysis pipelines, there are many elements to the pipeline described here. Elements of the pipeline can be partitioned separately and used as templates that establish a basis for mapping new analysis to previously benchmarked and verified templates. Here, we describe a set of 4 BCOs that closely mimic a real use case. The BCOs appear exactly as they would for a formal regulatory submission, and neatly capture several elements of the analysis in a way that brings clarity to an otherwise complex constellation of tools and pipelines. The BCOs clearly separate each element of the analysis in a way that is easy for a newcomer to follow, and easy to recapitulate in other analyses.

## Methods

All computations were carried out in HIVE (publicly available at https://hive.aws.biochemistry.gwu.edu/dna.cgi?cmd=home).

### Databases and Tools

#### Model Viruses and Host Genome

In order to represent a diversity of viruses, one virus was selected from each family listed in the human viruses page of the ViralZone knowledge base (https://viralzone.expasy.org/678) [35]. For those viruses with multiple NCBI RefSeq accessions, only one accession was used to generate synthetic reads. This process resulted in 25 reference genomes from 25 viruses that were used to synthetically spike the host read set (S1 Table).

#### Host Genome

To simulate the complexities of potential biological material in a synthetic read set, genetic sequence from other organisms was spiked into the data. The purpose was to plausibly simulate the presence of contaminating DNA originating from other steps in the vaccine production process. For this, chromosome Y was selected from the human genome to reflect the production of cell substrate from which a reagent is derived. The UniProt Proteome accession was selected for Human and the GenBank accession of chromosome Y was selected for analysis (S2 Table).

#### Curated viral database

The clustered version of the FDA reference viral genomes database (C-RVDB) v18 was downloaded from https://rvdb.dbi.udel.edu and processed using a curation method adapted by Lu’s and Salzberg’s method for removal of specific undesired sequences from genome databases [36]. Undesired sequences are considered those that best match non-viral sources, and therefore are not specific, or not uniquely specific, to known viruses. The resulting curated database, then, would be useful for detection of non-endogenous viruses, but not necessarily useful for evaluating endogenous retroviral or other retro-element sequences that exist within the non-viral animal genomes used in masking (Fig 1). Each of the viral genomes (n = 771,119) was split into 100 bp overlapping pseudo-reads using a sliding window of 50 bp, while respective coordinates and genome source information were recorded. The resultant pseudo-reads were taxonomically classified using Kraken2 [37] based on their alignment to a custom-built database of undesired sequences generated from the following NCBI (https://www.ncbi.nlm.nih.gov) records (Release 93, 2019): archaea, bacteria, plasmid, human hg38, fungi, plant, protozoa, UniVec_Core and coliphage phi-X174. DustMasker (part of the BLAST v2.3.0 release) (https://www.ncbi.nlm.nih.gov/books/NBK131777/) was employed to mask low complexity regions within the raw pseudo-reads. Finally, the resultant classified pseudo-reads from Kraken2 [37] and DustMasker[38] were used to mask the viral genomes database (Fig 1). Masking was done by converting the undesired sequence nucleotide letters to “N”. Importantly, to facilitate integration with HIVE we changed the format of headers (https://github.com/GW-HIVE/scripts) as well as avoided masking of the last 50 bp in each genome. The last 50 bp were masked only if these regions were classified as undesired. We also note that two copies of the genome NC_028379.1 was present in the C-RVDB (v18, n = 771,119 records), and only one copy was retained in the final curated database (CC-RVDB; n = 771,118 records).

**Fig 1.**
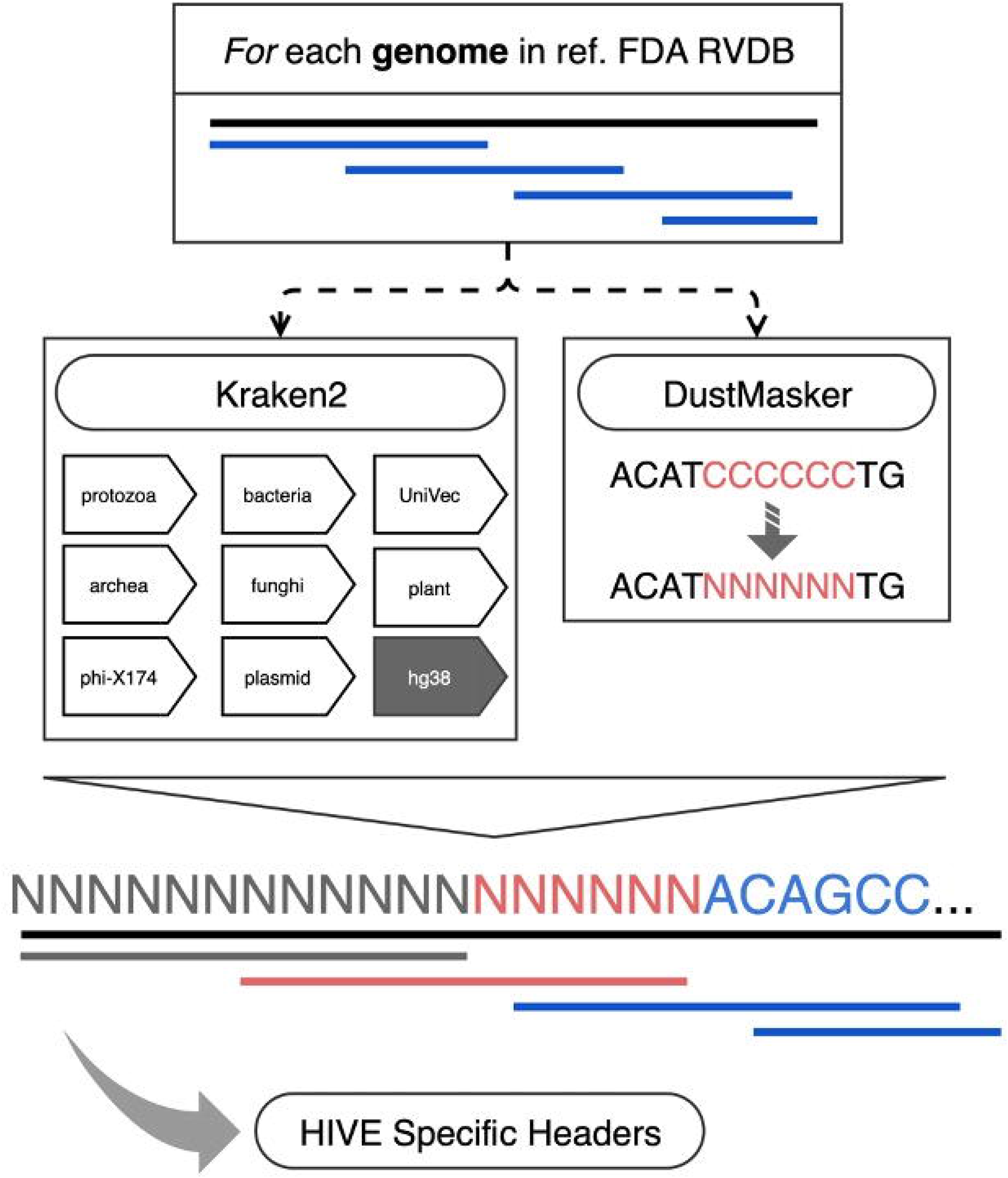
Viral genome database curation pipeline. (Top to bottom) Each genome within the FDA viral genomes database (black line, RVDB v18) was split into overlapping pseudo-reads (blue lines, 100 bp in length, with window size of 50 bp) that were systematically aligned to other target curation reference databases (as indicated in arrows) using Kraken2 (default setting, databases as indicated). Pseudo-reads were also scanned for low complexity regions using the DustMasker. Positives hits from these analyses were used to generate a curated version of the viral database that also includes HIVE-specific headers format for seamless integration with HIVE.

The source code for the database curation pipeline is available at https://github.com/Merck/curation-open-source. Analysis of genomes masking revealed on average low masking per genome (mean value = 11.8%; median value = 3.9%; Fig 2), yet some viral genomes were completely masked (∼1%). Although overall masking was low, it had a profound impact on the overall false positive (non-viral or not uniquely viral) detection rate in spiked-in samples.

**Fig 2.**
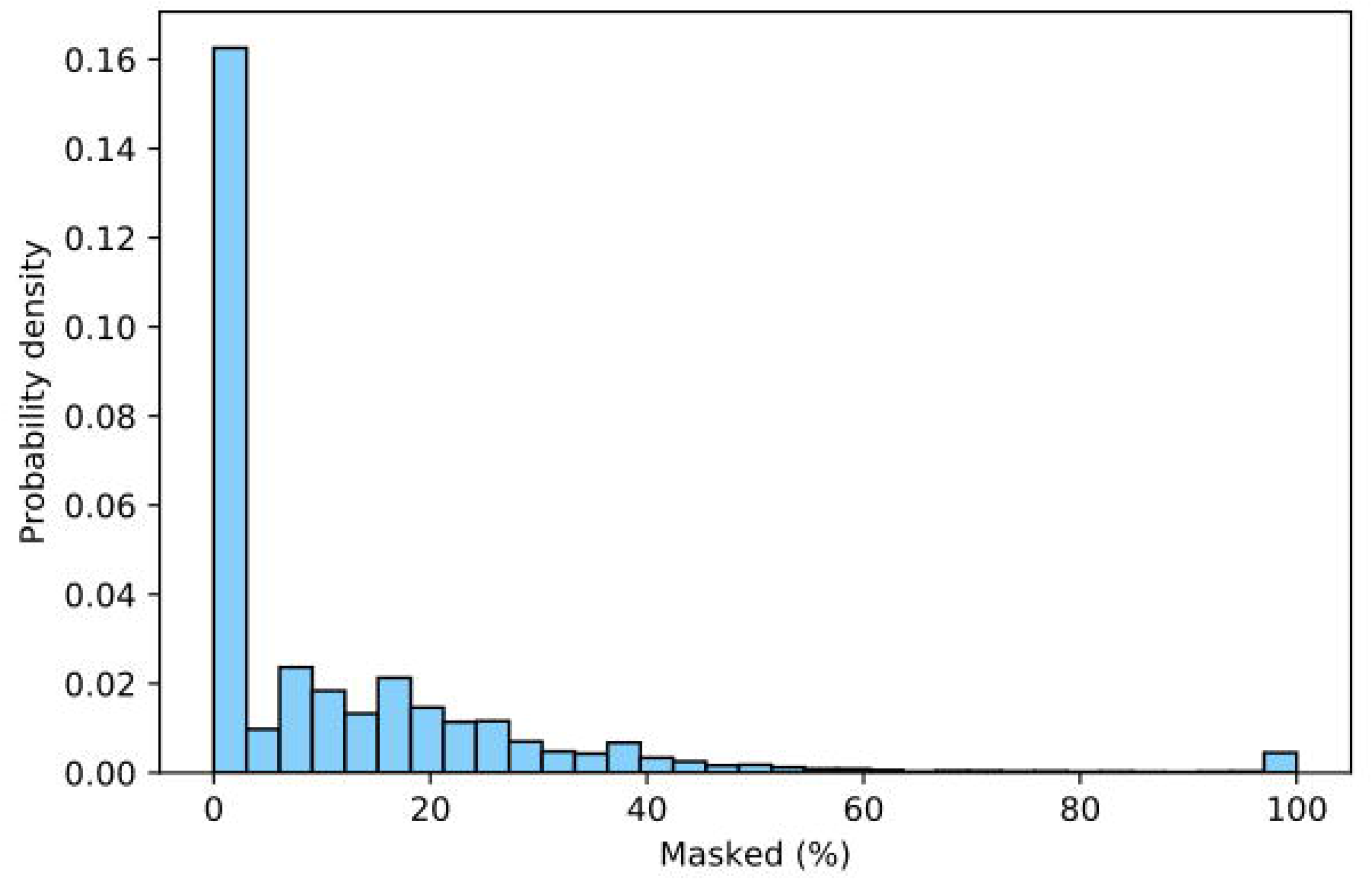
Distribution of percentage masking per viral genome within the FDA RVDB v18 viral genomes database. Each genome was split into overlapping pseudo-reads (100 bp in length, with a window size of 50 bp) that were systematically scanned for low complexity regions using DustMasker and aligned to target curation reference databases using Kraken2. Positives hits from these analyses were used to generate the final version of the curated database.

#### Generation of synthetic sample SN_25

Synthetic reads (paired-end) were generated from 25 model virus sequences (S1 Table) and Chromosome Y reference from *Homo sapiens* (CM000686.2) (S2 Table) to represent potential production-related cell substrates using the DNA *in*-*silico* tool [29]. One composite sample (SN_25), with a sample size of 63,667 reads, was created by mixing the synthetic reads from the 25 viruses in different ratios and 3,000 synthetic reads from Chromosome Y reference of *Homo sapiens* (CM000686.2), using the HIVE-seq tool [29]. The read length (range 35 to 150), the number of sequences from each reference and the average quality of the samples is described in S3 Table. The highest number of reads (10,000) were generated from *European bat lyssavirus* (NC_009527), *Banna virus* (NC_004211), *Adeno*-*associated virus* (NC_001401), *Varicella*-*zoster virus* (NC_001348), *Sapporo virus* (NC_006554), and *West Nile virus* (NC_001563) and the lowest number of reads (one) was generated from the *Influenza B virus* (NC_002204), *Human immunodeficiency virus* (NC_001802), *Hepatitis delta virus* (NC_001653), *Human Astrovirus* (NC_001943), *Poliovirus* (NC_002058), *Human Adenovirus* (NC_001405.1), and *SARS*-*CoV*-*2* (MT049951.1). The goal was to generate a heterogeneous sample of the known number of sequences to validate the parameter settings of the virus detection algorithm. All other general and advanced parameters used in the DNA *in*-*silico* tool were set to default.

#### Algorithm for virus detection in HIVE

The adventitious virus detection pipeline was developed in HIVE [29], [28] using previously described tools for the analysis of data and visualization of results. The workflow is illustrated in Fig 3 and explained in following steps.

**Fig 3:**
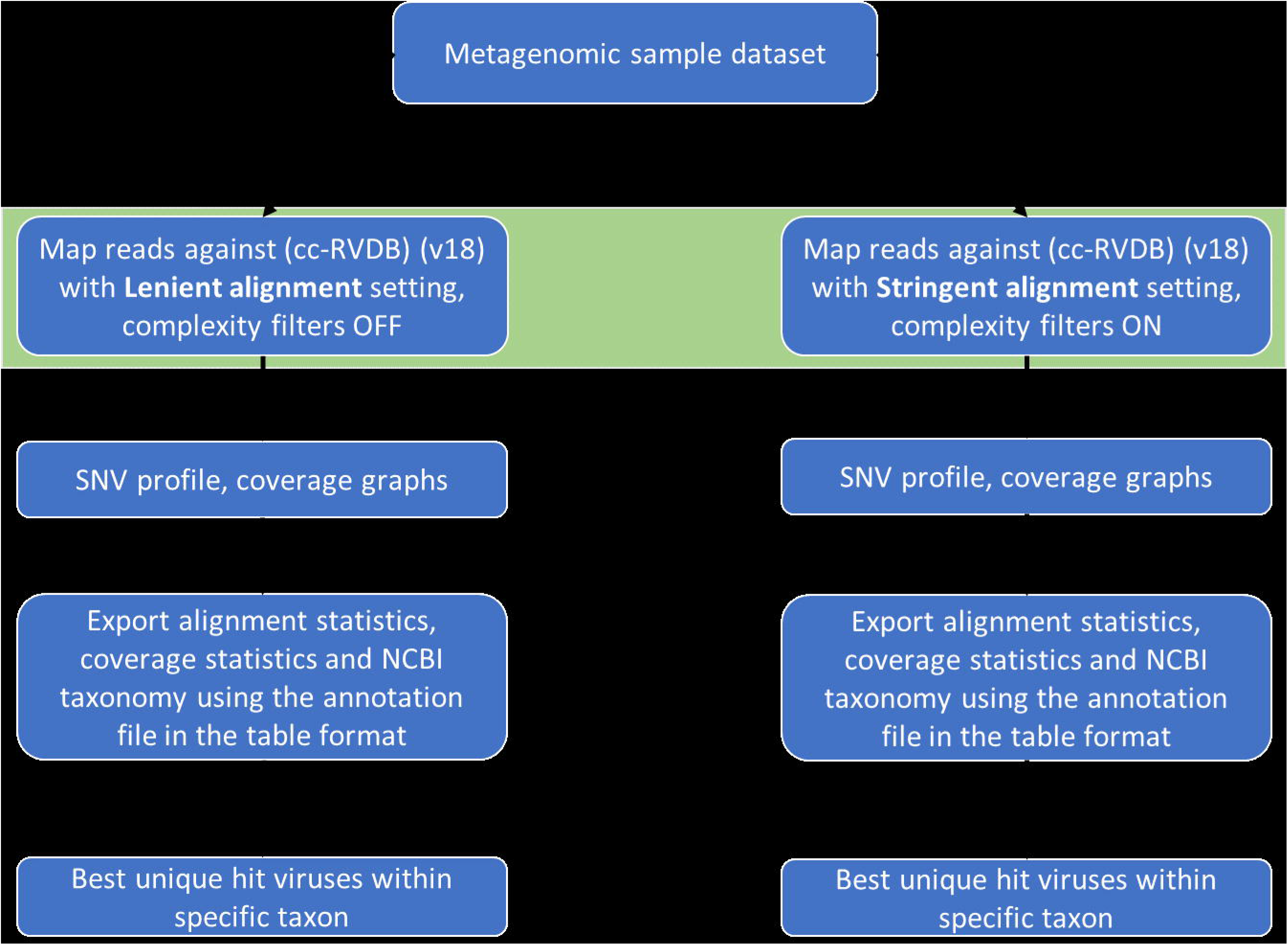
Simplified view of the analysis workflow. The lenient and the stringent pipelines, respectively are run in parallel and include four steps, which differ only in their alignment settings and complexity filters. In the lenient pipeline SN_25 is mapped against the CC-RVDB (v18.0) using HIVE-Hexagon aligner under lenient conditions with complexity filters turned off while as in the stringent pipeline alignment settings are strict and complexity filters are turned on. The SNP profiler is then run on the alignments to generate the respective coverage statistics. Coverage statistics and NCBI taxonomy for each hit (accession) are exported in extended tables using the annotation file for CC-RVDB (v18.0). Finally, best hits are evaluated manually to determine the accession with maximum hits or greatest total bases covered (total contig length) within each desired taxon.

### Analysis pipelines

For the sake of illustration, two analysis pipelines were used. These differed only in the initial alignment settings using the HIVE-Hexagon aligner but were identical in subsequent processing steps. Alignment settings were designed to illustrate the effect of either greater or lesser alignment stringency on the number of unique organisms hit in the viral database, some of which would be considered false positive hits. For example, the shorter minimum match lengths and higher allowed mismatches might be expected to result in hits to motifs that might be shared across many unrelated organisms. The Hexagon alignment settings that distinguish the lenient and stringent pipelines are documented in the Table 1.

**Table 1.** Hexagon alignment settings that distinguish the lenient and stringent pipelines. Three parameters intrinsic to the HIVE Hexagon tool are listed, along with their settings for either lenient or stringent alignments.

| Parameter | Lenient alignment | Stringent alignment |
| --- | --- | --- |
| Minimum Match Length (bases) | 50 | 75 |
| % mismatch allowed on alignment length | 15 | 5 |
| Complexity filters | Off | On (20 bases, "moderate" setting, on reads and references) |

The minimum match lengths were selected based on our anecdotal experience that shorter reads and more allowed mismatches result in more total hits. However, many of these have Total Coverage Lengths (TCL) that are too short to reliably indicate the presence of virus – they are arguably considered false positive hits or background noise. On the other hand, an overly stringent alignment may not detect potential viruses in a sample that deviates from the viral database or are novel and thus not well represented in the database’s viral sequence space. We do not claim that either of these pipelines is “optimized” for virus detection. However, they illustrate differences in performance and how settings are captured and distinguished in BCOs.

Other settings were common to both pipelines: seed Kmer=11, Markovnikov rule set. Also, in both pipelines, all equally best hits were counted. This choice has the advantage of producing the greatest TCL of any given accession but also results in total hits that are greater than the number of reads in the sample since some reads equally best hit multiple accessions (especially shorter reads where the higher proportion of mismatches are allowed).

Since the viral database was already curated against Human and vectors, the alignment of the sample, SN_25 (containing primarily non-viral background reads from human), was performed only against the curated version of the viral database CC-RVDB. In the event that a read set is known to contain a large proportion of reads attributable to a production cell species not included in the database curation in future experiments, it may be reasonable to include a concurrent counterscreen of reads against the CC-RVDB and the other cell species as long as the cell species genome is examined to assure that it does not contain viruses of interest in the overall detection study.

After the Hexagon alignment is completed, the Heptagon sequence profiling tool is run to compile variants and coverage statistics against the hit viral references from the CC-RVDB and generate graphical coverage profiles.

A tabular extended hit list is produced using an annotation file generated for the reference viral database, which guides the content in columns in the final extended hit list. It also contains the unparsed NCBI taxonomy for each accession in the database. The extended hit list compiles all accessions that were hit in the alignment along with their coverage statistics and unparsed NCBI taxonomies.

The final step in evaluating an alignment run is to determine the accession (and thus organism) with either most reads within each desired taxon or the greatest TCL. This is accomplished either by sorting the extended hit in order of either total hits or total contig length and secondarily by taxa. Such a sort is easily accomplished using a pivot table in Excel after parsing the NCBI taxonomy into separate columns, for instance. However, the present study focused only on a set of known viruses that were either endogenous or synthetic virus reads, so the results were obtained by simply sorting the extended hit list by the known taxa and selecting the accession with either greatest hits or greatest total contig length.

### Generation of BioCompute Object (BCO)

The steps of the pipeline, including the input data, metadata, and parameter settings, were documented as a BioCompute Object. HIVE’s built-in ability to record an analysis pipeline as a BCO was used to export analysis details for all computational steps, including the Description Domain, Execution Domain, IO Domain, and Parametric Domain. The automatically generated BCO was manually checked for accuracy, and information was added for the Provenance Domain and the Usability Domain, as well as content in the Description Domain and the “spec_version” field in the metadata. Quality checking steps were included in the Error Domain. No Extension Domain was used. Data was exported as a .json file. The BCO Portal (http://portal.aws.biochemistry.gwu.edu/) was used to generate the “etag” and “object_id.”

### Definition of terms

#### False positive

This term describes a hit in the reference database to accession or region within an accession that is arguably non-viral. Unless a concurrent “counterscreen” is performed (alignment against the viral database and cell-substrate genomes at the same time) or an initial cell-subtraction is performed (alignment against the cell-substrate genome first, and then alignment of cell-unaligned reads against viral database), some of the reads that hit an accession in the viral database may actually align as well or better to non-viral accessions in the broader GenBank NT. Examples might include ubiquitin sequences in some bovine viral diarrhea virus accessions or oncogenes in some retrovirus accessions; or integration region sequences; or even a viral sequence that is widely used in vectors and thus is not uniquely specific to the virus (cytomegalovirus (CMV) promoter, simian vacuolating virus 40 (SV40) terminator etc.). Database curation is intended to help reduce the opportunities for false positive hits. For purposes of virus detection in biological manufacturing processes for human or animal medicines, false positives increase the computational and manual work required to follow up categories of hits and thus reduce the efficiency of the tool for viral risk assessment or quality control.

#### False negative

This term can be applied to a viral read or population of reads that fail to hit a corresponding accession in the viral database. Failure to find a match in the viral database could represent a gap in the phylogenetic space covered by the viral database or could be attributable to overly stringent alignment settings. For purposes of virus detection in biological manufacturing processes for human or animal medicines, false negatives represent blind spots in the viral risk assessment.

#### Specificity

This term reflects the ability of the analysis pipeline to correctly attribute a read or population of reads to the truest matches and is influenced by the stringency of the alignment settings, the completeness, and curation of the viral database (or in lieu of curation, the counterscreen or cell-substrate subtraction process that might be applied).

#### Sensitivity

This term reflects the lowest level of specific reads or coverage that support the assertion that a specific virus is detected in the analysis pipeline. This is influenced by the alignment settings and processivity. Ideally, any specific read should find its best match, regardless of the amount of background non-specific reads. The decision on how much coverage, and whether it is based on a proportion of a genome or an absolute number of bases, of a given virus genome is required to assert specific detection also impacts sensitivity and may differ among different users or applications of the analysis pipeline.

#### Scalability

This term reflects the ability of the analysis pipeline and compute infrastructure to accommodate smaller (say <1 million) to much larger (say up to or >1 billion) populations reads in a given analysis or more complex computations such as translated alignments.

## Results

To more clearly delineate each component of the analysis, each of four workflows was packaged as a BCO, the results of which are described below. The workflow and result represented by each BCO are thereby more easily interrelated to each other conceptually as standalone units. Each of the four BCOs conform to IEEE2791-2020 and can be cross-referenced internally or by other BCOs, either as a citation upon which further work is built, or as a template for additional work.

The first BCO (S1 File) represents a command-line workflow for the curation of the viral database and was generated by manually entering steps into an online form-based editor (http://w3id.org/biocompute/portal, manuscript submitted). The BCO describes the entire pipeline and all inputs for a recipient or can be used to guide its re-execution in a step-by-step way, if the recipient has access to the data.

The other three BCOs (S2 File, S3 File, and S4 File) were generated using a built-in tool native to the HIVE platform. These BCOs describe the generation of synthetic reads for pipeline testing and the construction of the virus detection pipeline itself with lenient and stringent alignment settings to illustrate the comparison of results. A granular description of the results of these pipelines follows.

### Pipeline Performance

The performance of the two analysis pipelines was first evaluated using a contrived sample read set SN_25 composed of a known number (S3 Table) of *in silico* reads generated from 25 viruses (S1 Table), plus a background of 3,000 *in silico* reads generated from the Chromosome Y (CM000686.2) of the human genome. The evaluated results are documented in Table 2.

**Table 2.** Evaluation of the lenient and stringent alignment settings in virus detection. The sample SN_25 was generated by mixing the synthetic reads from 25 viruses and human Chromosome Y sequence in different proportions (S3 Table). The read length and different relative proportion of reads in SN_25 was documented in S2 Table. A comparison of the results in lenient and stringent alignment settings in virus detection pipelines is illustrated in the table. The second column documents the input reads from each virus in SN_25. The third and fourth columns document the best hit unique accession within a taxon and the maximum bases covered in the lenient and stringent pipelines, respectively. In general, from a sample size of 63667 reads, the lenient pipeline recovered 350 unique accessions with 51768 aligned reads, whereas stringent pipeline 219 unique accessions with total aligned reads 38061. Most of the reads that were not recovered by both the pipelines had read length shorter than the minimum match length threshold in lenient and stringent alignment. For example, JC polyomavirus with a max read length of 62 bases and Seoul virus with a max read length of 73 bases were filtered out in the Stringent alignment.

| Summary | Input reads | Lenient pipeline | Stringent pipeline |  |  |
| --- | --- | --- | --- | --- | --- |
| Total unique accessions hit (rows in extended hit list) |  | 350 | 219 |  |  |
| Total reads or reads aligned (hits) | 63,667 | 51768 | 38061 |  |  |
| Individual viruses (best hit within genus by TCL) | Input reads | Max of Hits (Max of TCL) | Max of Hits (Max of TCL) | Family | Genus |
| *Human adenovirus* | 1 | 1 (79) | 1 (79) | *Adenoviridae* | *Mastadenovirus* |
| *Human astrovirus* | 1 | 1 (79) | 1 (79) | *Astroviridae* | *Mamastrovirus* |
| *SARS-CoV-2* | 1 | 1 (79) | 1 (79) | *Coronaviridae* | *Betacoronavirus* |
| *Influenza B virus (RNA 1)* | 1 | 1 (79) | 1 (79) | *Orthomyxoviridae* | *Betainfluenzavirus* |
| *Poliovirus* | 1 | 1 (75) | 1 (75) | *Picornaviridae* | *Enterovirus* |
| *Human immunodeficiency virus* | 1 | 1 (79) | 1 (79) | *Retroviridae* | *Lentivirus* |
| *Hepatitis delta virus* | 1 | 1 (79) | 1 (79) | *Ribovirus* | *Deltavirus* |
| *Hepatitis B virus* | 10 | 1 (109) | 2 (193) | *Hepadnaviridae* | *Orthohepadnavirus* |
| *Bunyavirus La Crosse* | 10 | 3 (373) | 6 (736) | *Peribunyaviridae* | *Orthobunyavirus* |
| *Lake Victoria marburgvirus* | 10 | 2<br>(151) | 1<br>(95) | *Filoviridae* | *Marburgvirus* |
| *Human papillomavirus* | 10 | 6<br>(505) | 7<br>(605) | *Papillomaviridae* | *Alphapapillomavirus* |
| *JC polyomavirus* | 10 | 2<br>(111) |  | *Polyomaviridae* | *Betapolyomavirus* |
| *Mokola virus* | 10 | 8<br>(595) | 5<br>(436) | *Rhabdoviridae* | *Lyssavirus* |
| *Lassa virus* | 100 | 60<br>(3263) | 48<br>(3000) | *Arenaviridae* | *Mammarenavirus* |
| *Seoul virus* | 100 | 46<br>(1268) |  | *Hantaviridae* | *Orthohantavirus* |
| *Torque teno virus* | 100 | 38<br>(1879) | 21<br>(1317) | *Anelloviridae* | *Unclassified* |
| *Hendra virus* | 100 | 74<br>(5781) | 56<br>(4968) | *Paramyxoviridae* | *Henipavirus* |
| *Variola virus* | 100 | 43<br>(4129) | 34<br>(3524) | *Poxviridae* | *Orthopoxvirus* |
| *Mayaro virus* | 100 | 53<br>(3926) | 41<br>(3794) | *Togaviridae* | *Alphavirus* |
| *West Nile virus* | 10000 | 3186<br>(9985) | 2564<br>(9930) | *Flaviviridae* | *Flavivirus* |
| *Sapporo virus* | 10000 | 5253<br>(7474) | 6240<br>(7409) | *Caliciviridae* | *Sapovirus* |
| *Varicella-zoster virus* | 10000 | 6438<br>(102870) | 4579<br>(99376) | *Herpesviridae* | *Varicellovirus* |
| *Adeno-associated virus* | 10000 | 3862<br>(4050) | 3262<br>(4089) | *Parvoviridae* | *Dependoparvovirus* |
| *Banna virus (Segment 1)* | 10000 | 5677<br>(3743) | 5572<br>(3745) | *Reoviridae* | *Seadornavirus* |
| *European bat lyssavirus* | 10000 | 7599<br>(10910) | 5648<br>(10910) | *Rhabdoviridae* | *Lyssavirus* |

Out of 63,667 input sequences which includes 3,000 reads from host organism, 51,768 reads (81.3%) and 38,061 reads (59.8%) were recovered from the lenient and stringent pipelines, respectively. In general, the best hit accession for which results are recorded in the table did not recover all of the corresponding *in silico* reads for the respective virus. This is not surprising because some of these reads were filtered out based on their read lengths shorter than the threshold of 50 bases and 75 bases in the lenient and stringent pipelines, respectively. In addition, some reads may have best matched other accessions within a given taxon in the viral database, which ultimately were not the best or equally best hit documented above. Arguably, the sensitivity of the method could be improved by tallying all reads finding matches within the respective taxa rather than identifying just the single best hit or equally best hit accessions. JC polyomavirus and *Seoul virus* reads, with maximum read length of 62 bases and 73 bases, respectively (Table 2), were not recovered in the stringent pipeline because those reads did not qualify the minimum match length threshold set at 75 bases.

### Biological Data Test

Performance was also evaluated using a read set generated in a spiking study published in Dr. Arifa Khan’s paper “A Multicenter Study To Evaluate the Performance of High-Throughput Sequencing for Virus Detection” [39] and available publicly from the NCBI short read archive (Table 3). The sample was uploaded in HIVE, evaluated for quality using MultiQC tool[28], and analyzed using the lenient and stringent pipelines in the HIVE. The Illumina short read sample (SRX2804120) generated by LabB-2 represented HeLa cells spiked with a panel of known viruses: *Epstein-Barr virus* (EBV), *Respiratory syncytial virus* (RSV), *Feline leukemia virus* (FeLV), *Reovirus 1* (Reo1), and *Human papillomavirus 18* (HPV18). The average read length of the paired-end Illumina sample is 100 bases with average quality of reads as 33.9 and 35.4 Phred score, respectively. In both pipelines, the best hit accession (of an organism) within a specific taxon, as defined by the greatest number of bases covered (total coverage length), and TCL were captured. For EBV 1,216,173 and 672,477 unique hits within a taxon were retrieved in lenient and stringent alignments compared to 12 total hits in published results. Generally, the stringent pipeline was able to recover the accession with the greatest TCL for EBV, Reo1 and HPV18 compared to the published results. In both lenient and stringent pipelines, most of the hits retrieved for FeLV were for either oncoviruses or partial genomes except the three accessions, which do not contain arguably non-viral cellular sequence analogs, suggesting a comparatively low level of unique sequence among the FeLV reads. Some of the accessions for HPV18 were partially hit by 1-3 reads and their coverage statistics were not generated.

**Table 3.** Evaluation of Biological sample.

| Sample | Virus | Published results:<br>reads<br>(% coverage) | Lenient alignment:<br>reads (hits)<br>(% coverage) | Stringent alignment:<br>reads (hits)<br>(% coverage) | Family<br>Genus<br>Species |
| --- | --- | --- | --- | --- | --- |
| LabB-2<br>SRX2804120<br>Illumina short<br>reads | EBV | 13<br>(2.9) | 1216173<br>(54.6) | 672477<br>(68.4) | *Herpesviridae*<br>*Lymphocryptovirus*<br>*Human*<br>*gammaherpesvirus 4* |
|  | RSV | 1,176,557<br>(84.3) | 1351<br>(77.93) | 1808<br>(78.2) | *Pneumoviridae*<br>*Orthopneumovirus*<br>*Human*<br>*orthopneumovirus* |
|  | FeLV | 3,325<br>(94.7) | 5<br>(2.96) | 5<br>(5.3) | *Retroviridae*<br>*Gammaretrovirus*<br>*Feline leukemia virus* |
|  | Reo1* | 12<br>(10.4) | 5<br>(8.79) | 5<br>(12.71) | *Reoviridae*<br>*Orthoreovirus*<br>*Mammalian*<br>*orthoreovirus* |
|  | HPV18** | 3,800<br>(64.4) | 3000<br>(82.39) | 2582<br>(81.7) | *Papillomaviridae*<br>*Alphapapillomavirus*<br>*Alphapapillomavirus 7* |

The table documents the evaluation of the lenient and stringent alignment settings using the Illumina short reads generated in a spiking study representing the HeLa cells spiked with five model viruses Epstein-Barr virus, Respiratory syncytial virus, Feline leukemia virus, Reovirus 1, and Human polyoma virus18. The sample SRX2804120, publicly available from NCBI Short Read Archive (NCBI SRA), was uploaded in HIVE and evaluated for quality. The first three columns (1-3) represent the name of the sample, the name of the virus, and published results about the total number of reads recovered and the percentage of total bases covered. Columns 4 and 5 represent the evaluation of SRX2804120 using the lenient and stringent pipelines. The columns 4 and 5 capture the best accession hit within the taxon and the greatest number of bases covered (total contig length). In general, the stringent pipeline recovered the best hit with the most extended total contig length compared to the published results except in RSV and FeLV. In our analysis, most of the FeLV reads were for oncogenes or partial genomes, and some of the HPV18 accessions were hit partially by just 1-3 reads, and their coverage statistics were not generated.
\*Best hit accession in current results, not sum of all separate Reo1 genome segments
\*\*Not spiked; expected in HeLa cells

## Discussion

Until recently, communication of computational workflows in bioinformatics has been ad hoc. A biologist creates a mental checklist of things that she thinks are important for understanding the analysis, and communicates those elements in an impromptu way, without guidance for language or structure. This method of communicating a computational workflow is highly variable between researchers, and rarely sufficient to give the receiver full confidence or understanding of the pipeline, particularly for complex analyses that contain multiple parts.

Here, a suite of analyses is packaged as BCOs as they might appear in a formal regulatory submission. More specifically, a thorough comparison of lenient and stringent alignment settings in a virus detection analysis is presented, which includes the creation of a curated database and a synthetic read set. Submission-ready BCOs are generated for each major workflow for ease of comprehension and replication, and a bioinformatics platform on which the entire process is carried out is also described. The pipeline settings were initially tested and evaluated using a titration of the host background artificially spiked with synthetic reads and further demonstrated by analysis carried out with real biological data.

Every workflow and analysis necessary for understanding the process presented here is packaged as a BCO. Unlike existing workflow languages that focus predominantly on computational details, BCOs are report-ready formats that can be easily manipulated for ease of viewing. Because the underlying structure of a BCO is built on other computational standards like JSON Schema[40], they can also be programmatically interrogated or re-executed. The machine-readable format also makes them capable of integration with computationally-centric workflow languages that ease portability of execution, as described above, or for building customized views of the BCO report that present relevant content and collapse other details.

The four BCOs presented here represent different but related uses, each built with separate tools and with different focus. This presentation underscores the way in which an analysis can be compartmentalized into discrete modules that can be referenced in other contexts, either for reimplementation or reported through citation.

The first BCO describes the curation of the RVDB. The BCO was built manually using an online form-based editor, and reflects usage in a command line environment. Importantly, the BCO is used in the critically important step of database curation, which is often an overlooked and under described component of an analysis that can be difficult to understand or reproduce, but which influences all downstream steps. In this context, the database curation was important for generating higher confidence results with distinct identities in the database. The BCO unambiguously describes each step of the curation process.

This BCO also makes use of the optional “Extension Domain.” The Extension Domain is a user-defined domain that requires a schema to define its structure, and the contents of the domain. The Extension Domain is used to include additional structured information, if needed. To use the Extension Domain, a valid JSON schema for each extension used in this domain is expected to be specified through a link. The schema should be name spaced, and it is recommended that resolving the namespaced URI will provide the extension’s JSON Schema. The URL should be provided in the required “extension_schema” field. If greater execution portability is desired, then the included script can be in the Common Workflow Language v1.0 (https://w3id.org/cwl/v1.0/) or later format.

The first BCO is a data set BioCompute Object (dsBCO) and contains two different extension domains that follow this paradigm. A dsBCO is a computational record of a bioinformatics pipeline used to create a data resource. Objects such as this provide great clarity and transparency to the data integration processes. Both of the extension domains in the first BCO can be validated using the open source BCO-Tool v1.0.0 (https://github.com/hadleyking/bco-tool/tree/1.0.0).

The Source Configuration Management (SCM) extension (https://github.com/biocompute-objects/extension_domain/tree/1.1.0/scm) is used to record the location of source code for the BCO. The Additional Licenses and Dataset Categories (also known as “dataset_extension“) (https://github.com/biocompute-objects/extension_domain/tree/1.1.0/dataset) provides a vehicle for including licenses specific for the script and dataset created by the script. It also includes a set of tags for classification of the datasets.

The second BCO describes a process for generating synthetic “reads” from reference genomes as part of a common practice for testing the limits of a novel pipeline. The reads are generated by sampling input genomes from 25 viruses and Chromosome Y of human genome with known read lengths into one sample, SN_25, enabling precise measurement of pipeline performance for quality assessment.

The third BCO describes the virus detection pipeline using lenient alignment settings to recover unique organism hits, which uses both of the first two BCOs. The pipeline makes use of the curated database, which is the product of the first BCO and is tested using the synthetic reads generated in the second BCO. The pipeline itself describes the consequences of the lenient alignment settings in the viral detection using a database of known viral sequences, as is used in adventitious agent testing.

The fourth BCO describes the virus detection pipeline using stringent alignment settings to illustrate the difference in the detection of the unique virus hits compared to the lenient alignment settings. All other components of the pipeline are similar to the third BCO.

Although not specifically benchmarked, the utility of the virus detection algorithm in HIVE is comparable to existing bioinformatics pipelines such as SURPI [41] and VIP [42], with the advantage of very few non-specific hits with zero false positives in the results. The quality of results was further complemented by use of curated reference virus database [12]. The coverage graphs generated by the SNP profiler tool for each alignment and the user-friendly interface also made analysis easier and provided useful insights about the results. The “Extended Hit List” feature also provided granular details about the lineage information up to species taxonomy level, total contig length, percent mapped coverage per reference, and average coverage of contigs, adding an extensible element. This information is valuable in detection of any spurious hits in the results.

The last three BCOs describe the computational pipelines carried out on the HIVE platform. Both BCOs use the HIVE-native tool for BCO creation, which automatically pulls all relevant computational information and populates it in a BCO. File references are executed using URIs that reference a specific object ID, a native mechanism in HIVE, and similar mechanisms exist for other platforms. A user is prompted to order the steps, and then for metadata (e.g. Usability Domain and Provenance Domain).

The BCOs in this analysis describe the microcosm of interconnected analyses that depend on each other, and are therefore much more easily understood in a modular mechanism like BioCompute Objects. The constellation of interconnected tools and analyses can grow substantially larger, and BCOs are particularly helpful for quickly narrowing down to the precise element of the analysis in question, and the precise piece of information within that analysis, without presenting non-relevant information, but without discarding any of it – all of which can be very helpful for re-execution of a pipeline, and for asking targeted questions about intermediate files (e.g. .stat files that accompany many pipeline steps).

BCOs are currently accepted for regulatory submissions to the FDA for four drug applications by three Centers [18]. The small BCO files might in the future be easily appended to a formal regulatory submission as part of their own mechanism, such as by association in a Drug Master File (DMF) Type V. In this capacity, they can act as a manifest for all associated data files necessary for running or understanding the analysis, and recipe card for making sense of it all. A formal mechanism like this can help to improve the efficiency of workflow communication in a comprehensible way without compromising incoming data, which can still come in through traditional means. In addition, the integrity of the BCO itself can be checked for authenticity through the intrinsic “etag” [43], and can also be encrypted. Because the data is not stored on board with a BCO, the BCO is an ideal vehicle for a template format. The metadata focus can be very useful in a reporting context, such as when there is a need to explain how data is used (such as for transparency, reporting, or for critical review purposes) without disclosing the data, when there is a need (such as by regulators or reviewers) to target comments or questions to very specific aspects of a step within the protocol (or identify individuals associated with that step), or when there is a need for a data manifest to identify all incoming files, for example.

BioCompute can also support templating. Much scientific effort can be invested in developing, testing, and refining a workflow, and that results of those efforts can be captured in a template BCO (tBCO). The template can be referenced in future implementations of Run BCOs (rBCOs). Like platform specific templates for automation, the template could enable easy reimplementation, but in addition, a tBCO enables a researcher or regulatory reviewer to quickly become familiar with a template pipeline and check the validity of an instance against it.

In a regulatory environment, tBCOs could enable even more efficient communication of methods. A tBCO could be validated internally by the recipient, and checked against rBCOs for specific use cases. In this paradigm, the tBCO includes input data and an Error Domain (limits of detection of the pipeline, such as false negative rate, false positive rate, minimum read depth, or other relevant metrics). The input data and Error Domain are used to generate the “Validation Kit.” The Validation Kit evaluates the error in the pipeline, given some input. When a specific implementation of the pipelines is run as an rBCO, the output files are easily checked for validity against the Validation Kit to ensure that they fall within acceptable limits.

Ultimately, we feel that BCO usage, as described here, represents a substantial improvement on computational analysis communication. The important features of the BioCompute standard are that it provide enough structure to be useful for a recipient to be able to know what information to expect and where to find it with minimal effort, but that it also be flexible enough to accommodate new innovations in computational analyses and not inadvertently enforce any particular method for carrying out analyses. The BCO is platform agnostic, and has a heavy emphasis on human readability, acting more like a citation standard. The categorical partition of conceptually related information into top level “Domains” makes BCO navigation intuitive. Information about what parameters were used can be found in the Parametric Domain, for example. Finally, the flexibility of its machine readability makes viewing options for reviewers very easy, and is easily compatible with other computational mechanisms, such as scripts for re-execution. In total, we feel that the BioCompute mechanism represents at present the most straightforward and efficient way to communicate computational analyses, and which is not platform-dependent.

## Abbreviations

BCO: BioCompute Object
DMF: Drug Master File
FISMA: Federal Information Security Management Act
HIVE: High-performance Integrated Virtual Environment
IEEE: Institute for Electrical and Electronics Engineers
rBCO: runBCO
RVDB: Reference Viral Genomes Database
SCM: Source Configuration Management
tBCO: templateBCO
TCL: Total Coverage Lengths

## Funding Statement

This work was supported in part by Merck & Co., Inc., Kenilworth, NJ, USA, and by The McCormick Genomic and Proteomic Center at George Washington University

## Competing Interest Statement

Authors on this paper are/were employees of Merck Sharp & Dohme Corp., a subsidiary of Merck & Co., Inc., Kenilworth, NJ, USA and may hold stocks and/or stock options in Merck & Co., Inc., Kenilworth, NJ, USA.

## Supporting Information

**S1 Table. Viruses used to generate synthetic reads.** Each model virus used to generate synthetic reads, the family that it belongs to, and its accession number are listed.

**S2 Table. DNA sequences used to generate contaminating material.** Organism name and accession numbers are listed for each source of contaminant DNA.

**S3 Table. Composition of sample SN_25.** Sample SN_25 was generated as a composite of samples SN_1 - SN_24. The heterogeneous sample consisted of varying amounts of model virus in each of 6 different hosts (*Sus scrofa*, *Bos taurus*, *Gallus gallus*, Rhesus Macaque, *Homo sapiens*, and Chinese hamster), such that all samples are represented in varying amounts.

**S1 File. BCO describing the process for curating the FDA’s RVDB.** Description is provided for each of the steps, pipeline purpose and any idiosyncrasies, inputs, outputs, parameters, dependencies, prerequisites, execution environment, and potential error. Attribution, detailed data provenance are also provided. File may be more easily viewed in a JSON tree viewer or other visual-friendly online parser.

**S2 File. BCO describing the generation of 25 model virus samples.** Parameters and other relevant information (including unaligned statistics in the Error Domain as a means of estimating pipeline confidence) used in the generation of SN_1 - SN_25 are provided, along with attribution and data provenance.

**S3 File. BCO describing lenient alignment settings.** Parameters and other relevant information are described for the alignment procedure that used relaxed alignment criteria.

**S4 File. BCO describing stringent alignment settings.** Parameters and other relevant information are described for the alignment procedure that used strict alignment criteria.

## Notes

### Competing Interest Statement

The authors have declared no competing interest.

