## Supplementarty File 4 for "BioCompute Objects to communicate a viral detection pipeline with potential for use in a regulatory environment"

{
	"object_id": "https://w3id.org/biocompute/portal/BCO_00026693",
	"spec_version": "https://w3id.org/ieee/ieee-2791-schema/",
	"etag": "1b2b4e6c3475c0e4a6fbda88ab4d5362efa0ed789ca1d1e8f057d4bf8ed9fc9a",
	"description_domain": {
		"keywords": [
			"virus, metagenomics"
		],
		"xref": [
			{
				"namespace": "refseq",
				"name": "RefSeq",
				"ids": [
					"NC_001405",
					"NC_001563",
					"NC_003977",
					"NC_004297",
					"NC_001943",
					"NC_004108",
					"NC_005236",
					"NC_006554",
					"NC_045512.2",
					"NC_001608",
					"NC_002076",
					"NC_001348",
					"NC_002204",
					"NC_001352",
					"NC_001906",
					"NC_001401",
					"NC_002058",
					"NC_001699",
					"NC_001611",
					"NC_004211",
					"NC_001802",
					"NC_006429",
					"NC_009527",
					"NC_003417",
					"NC_001653"
				],
				"access_time": "2020-05-20T17:59:49-0400"
			}
		],
		"platform": [
			"HIVE"
		],
		"pipeline_steps": [
			{
				"step_number": 1,
				"name": "HIVE-Hexagon",
				"description": "HIVE hexagon alignment engine optimized for HIVE cloud performance on Next Generation data",
				"input_list": [
					{
						"uri": "https://hive2.biochemistry.gwu.edu/dna.cgi?cmd=objFile&ids=29990",
						"filename": "HIVE_C-RVDBv18.0_curated.fasta"
					},
					{
						"uri": "https://hive2.biochemistry.gwu.edu/dna.cgi?cmd=objFile&ids=32243",
						"filename": "SN_25"
					}
				],
				"output_list": [
					{
						"uri": "https://hive2.biochemistry.gwu.edu/dna.cgi?cmd=dna-hexagon&cmdMode=hexagon-&id=32247",
						"filename": "Strict_SN_25_new.fastq versus HIVE_C-RVDBv18.0_curated.fasta"
					}
				]
			},
			{
				"step_number": 2,
				"name": "Sequence Profiling Engine",
				"description": "Engine to perform coverage analysis ,base-calling, SNP calling ",
				"input_list": [
					{
						"uri": "https://hive2.biochemistry.gwu.edu/dna.cgi?cmd=dna-hexagon&cmdMode=hexagon-&id=32245",
						"filename": "Strict_SN_25_new.fastq versus HIVE_C-RVDBv18.0_curated.fasta"
					}
				],
				"output_list": [
					{
						"uri": "https://hive2.biochemistry.gwu.edu/dna.cgi?cmd=dna-heptagon&parent_proc_ids=32245&id=32247",
						"filename": "Strict_SN_25_new.fastq versus HIVE_C-RVDBv18.0_curated.fasta"
					}
				]
			},
			{
				"step_number": 3,
				"name": "Extended Hit List ",
				"description": "A tabular extended hit list is produced using an annotation file generated for the reference viral database, which guides the content in columns in the final extended hit list. It also contains the unparse NCBI taxonomy for each accession in the database. The extended hit list compiles all accessions that were hit in the alignment along with their coverage statistics and unparsed NCBI taxonomies.",
				"input_list": [
					{
						"uri": "https://hive2.biochemistry.gwu.edu/dna.cgi?cmd=dna-hexagon&cmdMode=hexagon-&id=32245",
						"filename": "Strict_SN_25_new.fastq versus HIVE_C-RVDBv18.0_curated.fasta"
					},
					{
						"uri": "https://hive2.biochemistry.gwu.edu/dna.cgi?cmd=dna-heptagon&parent_proc_ids=32245&id=32247",
						"filename": "Strict_SN_25_new.fastq versus HIVE_C-RVDBv18.0_curated.fasta"
					},
					{
						"uri": "https://hive2.biochemistry.gwu.edu/dna.cgi?cmd=record&ids=21220&types=u-ionAnnot",
						"filename": "RVDB_10June2020Update_Merged.tsv"
					}
				],
				"output_list": [
					{
						"filename": "o32247-reference612408-ProfileI",
						"uri": "https://hive2.biochemistry.gwu.edu/dna.cgi?cmd=dna-heptagon&parent_proc_ids=32245&id=32247"
					}
				]
			}
		]
	},
	"execution_domain": {
		"script": [
			{
				"uri": {
					"uri": "https://hive2.biochemistry.gwu.edu/dna.cgi?cmd=login"
				}
			}
		],
		"script_driver": "shell",
		"software_prerequisites": [
			{
				"name": "HIVE-Hexagon",
				"version": "1",
				"uri": {
					"uri": "https://hive2.biochemistry.gwu.edu/dna.cgi?cmd=dna-hexagon"
				}
			},
			{
				"uri": {
					"uri": "https://hive2.biochemistry.gwu.edu/dna.cgi?cmd=dna-heptagon"
				},
				"name": "Sequence Profiling Engine",
				"version": "1"
			}
		],
		"external_data_endpoints": [
			{
				"name": "HIVE",
				"url": "https://hive2.biochemistry.gwu.edu/dna.cgi?cmd=login"
			}
		],
		"environment_variables": {}
	},
	"io_domain": {
		"input_subdomain": [
			{
				"uri": {
					"uri": "https://hive2.biochemistry.gwu.edu/dna.cgi?cmd=objFile&ids=29990",
					"filename": "HIVE_C-RVDBv18.0_curated.fasta"
				}
			},
			{
				"uri": {
					"uri": "https://hive2.biochemistry.gwu.edu/dna.cgi?cmd=objFile&ids=32243",
					"filename": "SN_25"
				}
			}
		],
		"output_subdomain": [
			{
				"mediatype": "application/octet-stream",
				"uri": {
					"uri": "https://hive2.biochemistry.gwu.edu/dna.cgi?cmd=dna-heptagon&parent_proc_ids=32245&id=32247",
					"filename": "Strict_SN_25_new.fastq versus HIVE_C-RVDBv18.0_curated.fasta"
				}
			},
			{
				"uri": {
					"uri": "https://hive2.biochemistry.gwu.edu/dna.cgi?cmd=dna-heptagon&parent_proc_ids=32245&id=32247",
					"filename": "o32247-reference612408-ProfileInfo.csv"
				},
				"mediatype": "txt.csv"
			}
		]
	},
	"provenance_domain": {
		"name": "Viral metagenomics analysis using the stringent alignment settings",
		"version": "1.0",
		"review": [
			{
				"reviewer": {
					"name": "Naila Gulzar",
					"affiliation": "The George Washington University",
					"contribution": [
						"authoredBy"
					],
					"orcid": "https://orcid.org/0000-0002-8138-8312"
				},
				"reviewer_comment": "",
				"status": "approved"
			},
			{
				"reviewer": {
					"name": "Charles Hadley King",
					"affiliation": "George Washington University",
					"contribution": [
						"curatedBy"
					],
					"orcid": "https://orcid.org/0000-0003-1409-4549"
				},
				"status": "in-review",
				"date": "2021-05-03T20:19:00.000Z",
				"reviewer_comment": "Curated for BCO compliance"
			}
		],
		"contributors": [
			{
				"name": "Naila Gulzar",
				"affiliation": "The George Washington University",
				"contribution": [
					"authoredBy"
				],
				"orcid": "https://orcid.org/0000-0002-8138-8312"
			},
			{
				"name": "Jonathon Keeney",
				"affiliation": "The George Washington University",
				"contribution": [
					"authoredBy"
				],
				"orcid": "https://orcid.org/0000-0002-7665-6204"
			},
			{
				"name": "Jack Baker",
				"affiliation": "Merck & Co.",
				"contribution": [
					"authoredBy"
				]
			},
			{
				"name": "Geoffrey Hannigan",
				"affiliation": "Exploratory Science Center, Merck & Co.",
				"contribution": [
					"authoredBy"
				]
			},
			{
				"name": "Danny A. Bitton",
				"affiliation": "R&D Informatics Solutions, MSD Czech Republic s.r.o.",
				"contribution": [
					"authoredBy"
				]
			},
			{
				"name": "Ondrej Klempir",
				"affiliation": "Merck & Co.",
				"contribution": [
					"authoredBy"
				]
			},
			{
				"name": "Julia M Maritz",
				"affiliation": "Exploratory Science Center, Merck & Co.",
				"contribution": [
					"authoredBy"
				]
			},
			{
				"name": "Paul Duncan",
				"affiliation": "Merck & Co.",
				"contribution": [
					"authoredBy"
				]
			},
			{
				"name": "Raja Mazumder",
				"affiliation": "The George Washington University",
				"contribution": [
					"authoredBy"
				]
			}
		],
		"license": "https://spdx.org/licenses/CC-BY-4.0.html",
		"created": "2021-05-04T11:57:08.772Z",
		"modified": "2021-05-19T02:12:50.789Z"
	},
	"parametric_domain": [
		{
			"param": "alignmentEngine",
			"value": "1",
			"step": "1"
		},
		{
			"param": "keepAllMatches",
			"value": "3",
			"step": "1"
		},
		{
			"param": "minMatch",
			"value": "{29.0:{minMatchLen:75,minMatchUnit:0}}",
			"step": "1"
		},
		{
			"param": "mismatches",
			"value": "{30.0:{maxMissQueryPercent:5,considerGoodSubalignments:0}}",
			"step": "1"
		},
		{
			"param": "readconflicts",
			"value": "{resolveConfictsUnique:false,resolveConflicts:1,resolveConflictsScore:1}",
			"step": "1"
		},
		{
			"param": "complexity",
			"value": "{complexityQry:{11.0.0:{complexityEntropy:1.2,complexityWindow:20,maximumPercentLowQualityAllowed:0}},complexityRef:{11.1.0:{complexityRefEntropy:1.2,complexityRefWindow:20,acceptNNNQuaTrheshold:1}}}",
			"step": "1"
		},
		{
			"param": "name",
			"value": "Strict_SN_25_new.fastq versus HIVE_C-RVDBv18.0_curated.fasta",
			"step": "1"
		},
		{
			"param": "_type",
			"value": "svc-profiler-heptagon",
			"step": "2"
		},
		{
			"param": "name",
			"value": "Profile based on alignment : Strict_SN_25_new.fastq versus HIVE_C-RVDBv18.0_curated.fasta",
			"step": "2"
		},
		{
			"param": "directionalityInfo",
			"value": "true",
			"step": "2"
		},
		{
			"param": "generateIon",
			"value": "true",
			"step": "2"
		},
		{
			"param": "histograms",
			"value": "1",
			"step": "2"
		},
		{
			"param": "minCover",
			"value": "1",
			"step": "2"
		}
	],
	"usability_domain": [
		"Evaluation of the effect of the stringent alignment settings in the virus detection pipeline using the synthetic read set is demonstrated in this BCO. The alignment settings in HIVE-hexagon were designed to illustrate the effect of greater alignment stringency on the number of unique organisms hit in the viral database. An overly stringent alignment may not detect potential viruses in a sample that deviates from the viral database or are novel. The pipeline is comprised of an alignment, variant calling, and annotation generating steps. After the final computational step (step #3), a tabular extended hit list is produced using an annotation file generated for the reference viral database, which guides the content in columns in the final extended hit list. It also contains the unparse NCBI taxonomy for each accession in the database. The extended hit list compiles all accessions that were hit in the alignment along with their coverage statistics and unparsed NCBI taxonomies. The final step in evaluating an alignment run is to determine the accession (and thus organism) with either most reads within each desired taxon or the greatest total bases covered (total contig length). The extended hit list is opened in excel and sorted in order of either total hits or total contig length and secondarily by taxa. The present study focused only on a set of known viruses that were either endogenous or spiked, so the results were obtained by simply sorting the extended hit list by the known taxa and selecting the accession with either greatest hits or greatest total contig length."
	],
	"error_domain": {
		"empirical_error": {
			"Description": "Strict alignment executed 10 times with the same parameters, using synthetically generated data set based on West Nile Virus genome. Results were aligned to West Nile Virus reference genome using 'strict' pipeline parameters.",
			"aligned_standard_deviation": 50.437,
			"aligned_mean": 6447.5,
			"unaligned_standard_deviation": 50.43725,
			"unaligned_mean": 3552.5
		},
		"algorithmic_error": {
			"Description": "No fuzzy or stochastic processes in any of the algorithms used."
		}
	}
}
